# Identification of a novel biological role of arenavirus NP exonuclease (exoN) activity in a safe Pichinde virus model

**DOI:** 10.1101/2025.09.09.675144

**Authors:** Brigitte H. Flannery, Qinfeng Huang, Hinh Ly, Yuying Liang

## Abstract

Arenaviruses such as Lassa virus (LASV) and Junin virus (JUNV) can cause lethal hemorrhagic fever diseases with no FDA-approved vaccines or effective therapeutics. The arenaviral nucleoprotein (NP) contains an exoribonuclease (exoN) domain that suppresses type I interferon (IFN-I) production by degrading pathogen-associated molecular-pattern (PAMP) RNAs. The biological roles of NP exoN during viral infection have not been well characterized for highly pathogenic arenaviruses, largely due to biosafety constraints and challenges in generating viable NP exoN mutants. We previously developed a recombinant tri-segmented viral vector (rP18tri) based on non-pathogenic Pichinde virus (PICV), which allows functional studies of transgenes during arenavirus infection in a BSL-2 setting. In this study, we introduced point mutations at individual catalytic residues to abolish PICV NP exoN activity and generated exoN-deficient rP18tri(NPm) reporter viruses. These mutants failed to replicate in IFN-competent A549 cells, accompanied by strong IFN-I induction. Expression of wild-type (WT) JUNV NP, but not its exoN-deficient mutant, in the rP18tri(NPm)-G backbone restored single-cycle infectivity and suppressed IFN-I induction. Furthermore, exoN-deficient viruses produced disproportionately high levels of defective virion particles relative to virion RNA from infected A549 cells, a defect that was rescued by expression of WT JUNV NP but not the exoN-deficient NP as a transgene. Together, these data demonstrate an essential role of NP exoN in IFN-I suppression and identify a novel biological function of NP exoN in promoting the production of infectious virion particles in IFN-competent cells.

**Importance:** Arenaviruses, such as Junin virus (JUNV), can cause severe viral hemorrhagic fever in humans, with no FDA-approved vaccines or effective therapeutics. Arenaviral nucleoprotein (NP) contains an exoribonuclease domain (exoN), which helps to evade the host immune system by blocking the production of type I interferons (IFNs). However, the biological role of NP exoN in arenavirus infection is not well understood. In this study, we used a recombinant, tri-segmented non-pathogenic Pichinde virus (PICV) with NP exoN mutations as a safe and infectious arenavirus vector to express JUNV NP as a transgene. We found that JUNV NP exoN plays a key role in suppressing IFN-I responses and discovered a previously unrecognized function of exoN in producing infectious virion particles in IFN-competent cells. These findings improve our understanding of arenavirus infection and identify NP exoN as a promising target for developing new antiviral therapies.

## INTRODUCTION

Mammarenaviruses (arenaviruses) are mostly carried in local rodent species and are phylogenetically and geographically categorized into two major groups (1). New World (NW) arenaviruses are found in the Americas, whereas Old World (OW) arenaviruses are found in Africa, Asia, and Europe (2). Arenaviruses are zoonotic viruses that can infect humans through inhalation of virus-containing aerosolized urine/feces or consumption of contaminated food or water (3, 4). Several OW and NW arenaviruses can cause severe and sometimes lethal diseases in humans, including Lassa virus (LASV) in West Africa, and Junin virus (JUNV) and Machupo virus (MACV) in South America. Arenavirus-caused hemorrhagic fevers show similar symptoms, which can include fever, lethargy, and myalgia in mild cases, with severe cases resulting in bleeding disorders, seizures, and shock with a high fatality rate (5–7). There are no US Food and Drug Administration (FDA)-approved vaccines or effective antiviral therapeutics against these highly pathogenic arenaviruses (4), which are classified as category A priority pathogens by the Centers for Disease Control and Prevention (CDC) due to their significant risk to national security and public health and require handling in biosafety level (BSL)-3 or BSL-4 facilities. A live attenuated JUNV vaccine Candid#1 is licensed for use in Argentina against JUNV-caused Argentine hemorrhagic fever, but is unlikely to be approved by the FDA due to safety concerns (8). Ribavirin has been used to treat Lassa fever but it is only effective if given early during the course of the infection and has severe side effects (9). These reasons emphasize the need for the development of effective vaccines and antiviral therapeutics for arenaviral infections.

Arenaviruses are enveloped RNA viruses with a bisegmented, ambisense genome, encoding four viral gene products: L polymerase, nucleoprotein NP, glycoprotein GPC, and matrix protein Z (1). Although the pathogenic mechanism has not been well-characterized, the hallmarks of arenavirus hemorrhagic fevers are a high viremia and a generalized immune suppression (2, 6). Evidence from clinical studies and non-human primate models suggests that severe LASV infections are associated with the lack of early induction of innate immunity and the lack of effective cytotoxic T-cell responses (10–13). A major arenaviral innate immune antagonist is the NP protein, which has been shown to strongly block the induction of type I interferons (IFN-Is) mainly through its exonuclease (exoN) activity. The C-terminal region of all arenavirus NP proteins contains a conserved DEDD(h) exoN domain, which preferentially degrades double-stranded (ds) RNA *in vitro* and is critical for IFN-I suppression via degradation of PAMP RNAs (14–16). Most studies have relied on NP overexpression in transfected cells (17, 18) and have been limited to only a few arenavirus reverse genetic systems. We have shown that recombinant Pichinde virus (PICV) mutants with exoN catalytic mutations grew to high titers in IFN-deficient Vero cells but were replication-defective in IFN-competent A549 cells with highly induced IFN-Is, and completely lost virulence in a guinea pig model (19). Similar findings were observed with recombinant LASV with mutations at and near the exoN catalytic residues (20, 21). Most arenaviruses, including PICV and LASV, do not activate IFN-Is upon infection in cell cultures (19, 20). Consistently, little to no IFN-I production has been detected in arenavirus animal models and in Lassa fever patients (13, 19, 22, 23). In contrast, JUNV infection was found to induce dsRNA accumulation and IFN-I production in cell cultures, consistent with high levels of serum IFN-I detected in severe Argentine hemorrhagic fever cases (19, 20, 23, 24), despite the fact that JUNV NP can strongly block IFN-I activation in plasmid-transfected cells and has a functional exoN domain (16, 25). Thus, whether JUNV NP exoN functions to suppress the IFN-I induction and what specific biological role(s) it plays during authentic arenaviral infection remain to be clarified. However, recombinant pathogenic arenaviruses with targeted exoN deficiency are difficult to generate and characterize because of the high-containment requirements and the multiple roles of NP in the viral life cycle. To overcome these limitations, our lab utilizes a tri-segmented recombinant PICV vector to safely study the biological roles of pathogenic arenavirus proteins during an authentic arenavirus infection at the BSL-2 level (26).

PICV is a non-pathogenic arenavirus isolated from rice rats in Colombia, South America (27). Sequential passage of PICV in guinea pigs led to a highly virulent strain (P18) that can cause a similar lethal disease in guinea pigs as human Lassa fever (28). As such, PICV infection of guinea pigs has been used as a safe surrogate model for Lassa fever (29, 30). We have established the reverse genetics system for PICV and developed a recombinant viral vector, rP18tri, by splitting the wild-type S segment into two, S1 and S2 segments, encoding essential viral genes GPC and NP on the S1 and S2, respectively, and an additional open-reading frame (ORF) on each of the segments (31). This rP18tri vector has been used to develop vaccine candidates with robust antibody and T-cell immunity (32–35). In addition, we used rP18tri to express immune antagonists, such as LASV Z, as a transgene and demonstrated the biological role of its RIG-i inhibition in arenavirus gene expression and viral infectivity (26). Expressing other arenavirus proteins as transgenes in the rP18tri vector system allows the separation of their essential roles in arenaviral replication from their biological functions in virus-host interactions.

In this study, we aimed to characterize the biological role(s) of JUNV NP exoN in the context of the rP18tri vector. We first introduced mutations to PICV NP exoN catalytic residues in the rP18tri-G reporter virus to eliminate the PICV NP exoN activity and cloned either the wild-type (WT) or exoN-deficient JUNV NP gene into the extra ORF of the rP18tri(NPm)-G vector. By comparing these recombinant viruses in IFN-deficient Vero cells and IFN-competent A549 cells, we confirmed the essential role of NP exoN activity in IFN-I suppression and arenavirus multiplication, and identified a novel biological role of NP exoN in promoting efficient production of infectious virion particles relative to virion RNA levels in IFN-competent cells.

## MATERIALS AND METHODS

### Ethics Statements

Research conducted for this manuscript was approved by the Institutional Biosafety Committee at the University of Minnesota, Twin Cities, under the protocol ID 2309-41371H.

### Plasmids, cells, and viruses

The three plasmids expressing the L, S1, and S2 RNA segments to generate recombinant tri-segmented PICV virus (rP18tri) have been described previously (31). To make exoN-deficient rP18tri(NPm) viruses, the PICV NP gene fragment with D389A (m1) or E391A (m2) mutation (19) was swapped into the S2 plasmid via conventional molecular cloning techniques. The wild-type (WT) JUNV NP gene (17) and a mutant carrying dual exoN catalytic mutations (D380N/E382Q) were cloned into the S1 plasmid in its extra open-reading frame (ORF). All plasmids were sequence-confirmed.

BHK-21 baby hamster kidney cells, African green monkey kidney Vero cells, and human lung epithelial A549 cells were maintained in Dulbecco’s modified Eagle’s medium (DMEM) supplemented with 10% fetal bovine serum (FBS) (Sigma) and 50 μg/ml penicillin-streptomycin (Invitrogen Life Technologies). BSRT7-5 cells, which are BHK-21 cells stably expressing T7 RNA polymerase, were obtained from K.K. Conzelmann (Ludwig-Maximilians-Universität, Germany) and were cultured in minimal essential medium (MEM) (Corning) supplemented with 10% FBS (Sigma) and 50 μg/ml penicillin-streptomycin (Invitrogen Life Technologies).

The tri-segmented rP18tri-based reporter viruses were amplified by infecting BHK-21 cells at a multiplicity of infection (MOI) of 0.01 for 48 hours. Viruses were quantified by plaque assay to determine plaque-forming units (PFU) per ml and by FACS to determine GFP-positive units (GPU) per ml.

### Generation of rP18tri-based reporter viruses expressing pathogenic arenavirus NP

Recombinant tri-segmented rP18tri-based reporter viruses were rescued as previously described (31). Briefly, the BSRT7-5 cells were transfected with three plasmids expressing the L, S1, and S2 RNA segments. The S1 plasmid contained the JUNV NP gene insert in the antisense ORF. The S2 plasmid contained wild-type (WT), D389A (m1), or E391A (m2) mutation in the PICV NP gene. Supernatants were collected from the transfected BSRT7-5 cells and analyzed by plaque assay on Vero cells. Recombinant viruses were collected from individual plaques and amplified on BHK-21 cells to generate recombinant virus stocks.

### Quantification of infectious reporter virus (GPU/ml) by flow cytometry (FACS)

Vero cells were seeded onto 12-well plates at 2x10^5^ cells/ml and infected with 250 µl of viral samples for one h at 37°C. Cells were collected for FACS analysis at 10 hours post-infection (hpi), which allows for the GFP expression during the first round of replication. Cells were trypsinized, washed with phosphate-buffered saline (PBS) (Sigma), then re-suspended in FACS buffer (PBS with 1% FBS) with 1% paraformaldehyde (Sigma Aldrich). Data acquisition was performed on BD FACSCelesta (BD Biosciences) to count 100,000 cells per condition. Data analysis was performed on FlowJo (BD Biosciences). The infectious virus titer was calculated by the number of GFP-positive cells, the infection volume, and dilutions to determine GFP-positive units (GPU) per ml.

### Quantification of GFP-positive cells by FACS

Vero or A549 cells were seeded onto 12-well plates at 2x10^5^ cells/ml and infected with virus at a designated MOI. Cells were trypsinized, washed with PBS (Sigma), then re-suspended in FACS buffer (PBS with 1% FBS) with 1% paraformaldehyde (Sigma Aldrich). Data acquisition was performed on BD FACSCelesta (BD Biosciences) to count 100,000 cells per condition. Data analysis was performed on FlowJo (BD Biosciences).

### Quantification of ISG expression by RT-qPCR

A549 cells were seeded onto 12-well plates at 2x10^5^ cells/mL and infected with WT and mutant rP18tri viruses at an MOI of 0.05. At 16 hpi, cell lysates were collected in 1X DNA/RNA Shield (Zymo Research). RNA was extracted from the cell lysates using the Zymo MagBead RNA Extraction Kit (Zymo Research). ISG expression was quantified by RT-qPCR using the Luna One-Step quantitative RT-qPCR kit (New England Biolabs) with gene-specific primers, 5’-CTCTGAGCATCCTGGTGAGGAA-3’ and 5’-AAGGTCAGCCAGAACAGGTCGT-3’ for ISG15, 5’-CGTGTGCCATTGTTCTCTTTG-3’ and 5’-GTCAGACTGCCTCATCTGTAAC-3’ for RIG-I, 5’-GAGGAATCAGCACGAGGAATAA-3’ and 5’-TCAGATGGTGGGCTTTGAC-3’ for MDA-5, 5’-GAAGTGGACCTCTACGCTTTGG-3’ and 5’-TGATGCCATCCCGTAGGTCTGT-3’ for PKR, and 5’-GAAGGTGAAGGTCGGAGTC-3’ and 5’-GAAGATGGTGATGGGATTTC-3’ for GAPDH. Each reaction was performed in technical duplicates and incubated at 55°C for 10 min, 95°C for one min, followed by 40 cycles of amplification at 95°C for 10 sec, 60°C for 30 sec, followed by a melting curve analysis. Fold changes in gene expression were determined by calculating 2^-ΔΔCq^ values using cellular Glyceraldehyde 3-Phosphate Dehydrogenase (GAPDH) as a reference gene.

### Quantification of IFN-β levels in supernatants by ELISA

A549 cells were seeded onto 12-well plates at 2x10^5^ cells/mL and infected with wild-type and mutant rP18tri viruses at an MOI of 0.05. Supernatant samples were collected at 16 hpi. The concentration of IFN-β in the supernatants was measured using the LEGEND MAX Human IFN-β ELISA kit (Biolegend), following the kit instructions. Briefly, samples or IFN-β standard were appropriately diluted, added to the plate pre-coated with human IFN-β antibody, and incubated overnight at 4°C with rocking. The plate was then washed, incubated with the human IFN-β detection antibody for 1 h, washed again, and incubated with the Streptavidin-Polymer HRP secondary antibody for 30 min. After a final wash step, detection reagents were added and the 450 nm absorbance was measured using a BioTek Synergy 2 plate reader (Agilent). IFN-β concentration was calculated using a linear regression of the standard curve in Prism (GraphPad).

### Quantification of viral RNA by RT-qPCR

Viral supernatants were filtered through 0.45 μm filters (Genesee Scientific) and subjected to RNA extraction using Zymo MagBead RNA extraction kit (Zymo Research). Serial 10-fold dilutions of WT PICV virion RNA were used to create a standard curve. RT-qPCR was performed using the Luna One-Step reaction mix (New England Biolabs) with primers 5’-CCCGGACAGAGAAATCCTTATG-3’ and 5’-CTCCCTTGAACTTGAGACCTTG-3’ specific for the PICV NP gene. Each reaction was performed in technical duplicate and incubated at 55°C for 10 min, 95°C for one min, followed by 40 cycles of amplification at 95°C for 10 s and 60°C for 30 s.

## RESULTS

### The tri-segmented rP18tri-G virus with NP exoN mutations replicated in IFN-deficient Vero but not in IFN-competent A549 cells

PICV NP exoN contains 5 catalytic residues D389, E391, D466, D533, and H528 (19). We have previously shown that mutation of each catalytic residue abolished the exoN enzymatic activity and NP-mediated IFN-I suppression and that recombinant PICV exoN mutants can only replicate in Vero but not in A549 cells, demonstrating an essential role of NP exoN for arenavirus replication in IFN-competent cells by inhibiting the IFN-I production (19). As the bi-segmented PICV genome cannot easily accommodate transgenes, we developed a tri-segmented PICV vector, rP18tri, and inserted the GFP reporter gene into the S2 segment (**Fig. 1A**, left). To eliminate the NP exoN activity in the rP18tri backbone, we introduced NP D389A (m1) or E391A (m2) mutation into the rP18tri-G genome and generated recombinant viruses (NPm1)-G and (NPm2)-G, respectively (**Fig. 1A**, right). While rP18tri-G infection led to GFP expression in both Vero and A549 cells, infection with (NPm1)-G and (NPm2)-G expressed GFP only in Vero but not in A549 cells at 48 hpi (**Fig. 1B**). These data are consistent with our previous study using the bi-segmented rPICV exoN mutants (19) and demonstrate that the NP exoN activity plays a critical role for viral replication in the IFN-competent cells possibly by blocking the IFN-I production.

**Figure 1.**
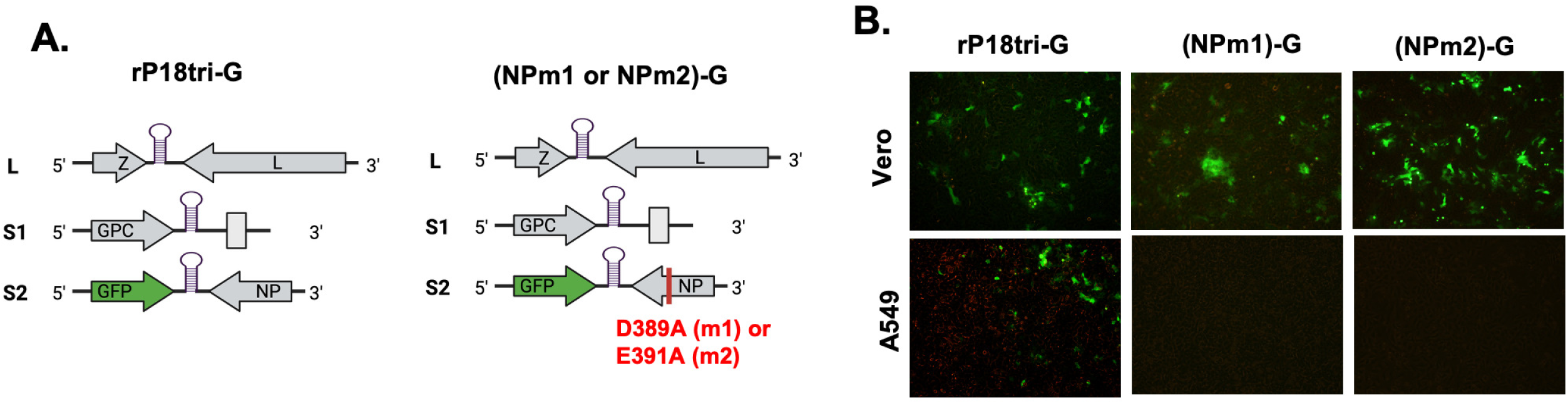
The tri-segmented NP exoN-deficient rP18tri-G mutants replicated in Vero but not in A549 cells. (**A**) Genome organization of rP18tri reporter virus with WT NP gene (rP18tri-G) (left panel) and NP exoN catalytic mutant D380A ((NPm1)-G) or E391A ((NPm2)-G) (right panel). (**B**) Infection of Vero and A549 cells by rP18tri-G, (NPm1)-G, or (NPm2)-G, each in triplicate. Representative GFP images at 48 hpi are shown.

### Expression of JUNV NP as a transgene did not rescue multiple-cycle viral infection of exoN-deficient rP18tri viruses in IFN-competent A549 cells

To determine whether expression of a functional arenavirus NP as a transgene can rescue the replication of rP18tri exoN mutants in IFN-competent cells, we cloned WT JUNV NP (JvNP) into the (NPm1)-G or (NPm2)-G virus genome backbone and generated the corresponding rP18tri virus (NPm1)-JvNP/G or (NPm2)-JvNP/G (**Fig. 2A**, top). As a control, an exoN-deficient JvNP (JvNPm) was cloned into the (NPm2)-G vector backbone to produce the recombinant virus (NPm2)-JvNPm/G (**Fig. 2A**, bottom).

**Figure 2.**
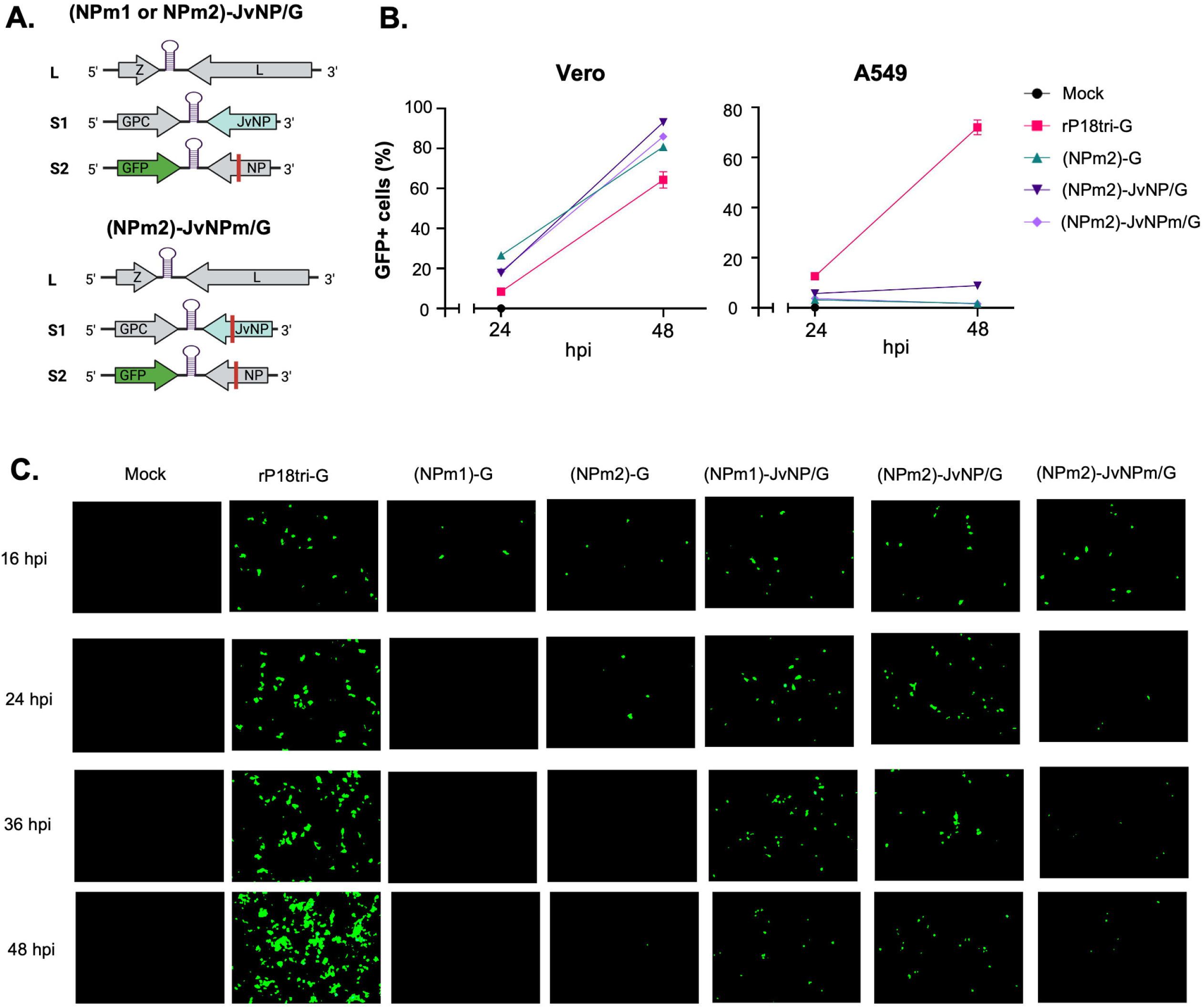
Expression of JUNV NP as a transgene did not rescue multi-cycle replication of exoN-deficient rP18tri-G mutants in A549 cells. **(A)** Genome organization of (NPm1 or NPm2)-JvNP/G and (NPm2)-JvNPm/G, which encode WT JUNV NP and exoN-deficient JUNV NP, respectively, in the negative sense on the S1 segment. (**B**) Viral growth in Vero and A549 cells. Cells were infected with mock or the indicated viruses, each in triplicate. The percentage of GFP-positive cells at 24 and 48 hpi was quantified by FACS and shown as an average with standard deviation for each virus. (**C**) Viral infectivity in A549 cells. Cells were infected with mock or the indicated viruses. Representative images of GFP expression at the indicated times post-infection are shown.

A set of (NPm2)-G viruses encoding an empty cassette, WT JvNP, and exoN-deficient JvNPm was used to infect Vero and A549 cells, along with mock and rP18tri-G controls. The percentage of GFP-positive cells, measured by FACS, increased from 24 to 48 hpi for all viruses in Vero cells, but only for rP18tri-G in A549 cells (**Fig. 2B**), suggesting that expression of WT JUNV NP as a transgene is unable to fully rescue the multiple-cycle replication of (NPm2)-G in IFN-competent cells.

To identify the stage(s) at which replication of these rP18tri exoN mutant viruses was inhibited in A549 cells, we monitored GFP expression in infected cells at 16, 24, 36, and 48 hpi using fluorescence microscopy (**Fig. 2C**). The control virus rP18tri-G led to increasing GFP expression over the time course, correlating with its efficient multiple-cycle replication in A549 cells as shown in Fig. 2B. For both (NPm1)-G and (NPm2)-G virus infections, GFP expression was detected in a small fraction of cells at 16 hpi but declined and disappeared thereafter, suggesting that these mutant viruses can enter cells and initiate viral transcription and translation, but their replication is aborted after the first cycle. For (NPm1)-JvNP/G and (NPm2)-JvNP/G viruses, which express WT JvNP as a transgene, GFP expression increased slightly from 16 to 24 hpi and decreased slightly by 48 hpi. In contrast, (NPm2)-JvNPm/G, which expresses an exoN-deficient JUNV NP, had similar GFP expression kinetics as (NPm1)-G and (NPm2)-G virus infections. These data demonstrate that expression of WT JUNV NP as a transgene is insufficient to fully restore viral multiplication of rP18tri exoN mutant viruses in IFN-competent cells, but JUNV NP exoN activity can partially restore viral infectivity during the early viral replication cycles.

### Expression of JUNV NP as a transgene rescued single-cycle viral replication of rP18tri exoN mutants in the IFN-competent A549 cells

To assess whether the differential GFP expression observed for infections of rP18tri exoN mutants in A549 cells (**Fig. 2C**) is independent of their intrinsic replication capacity, we infected Vero and A549 cells at a high MOI of 0.3, quantified GFP-positive cells at 16 hpi to assess single-cycle replication, and compared the relative infectivity between the two cell lines. Compared to (NPm1)-G infections, which resulted in moderate GFP expression (∼ 14%) in Vero cells but very low levels (∼ 1.3%) in A549 cells, (NPm1)-JvNP/G infections showed a moderate to high GFP expression (∼ 45% in Vero and ∼ 16% in A549) in both cell lines, similar to the control rP18tri-G virus (∼64% in Vero and ∼29% in A549) (**Fig. 3A**). The A549/Vero (A/V) ratio of GFP expression, representing the relative replication capacity in IFN-competent cells, ranges from 0.09 for (NPm1)-G) to 0.35 for (NPm1)-JvNP/G and 0.44 for rP18tri-G, suggesting that expression of JUNV NP as a transgene markedly increases viral infectivity of (NPm1)-G in the presence of IFN-Is. Similar results were obtained with (NPm2)-G, which showed significantly lower infectivity in A549 cells (A/V ratio = 0.07) compared to rP18tri-G (A/V ratio = 0.27) (**Fig. 3B**). (NPm2)-JvNP/G, which expresses WT JUNV NP as a transgene, showed infectivity (A/V ratio = 0.21) comparable to rP18tri-G, whereas (NPm2)-JvNPm/G, expressing an exoN-deficient JUNV NP, exhibited a similar infectivity (A/V ratio = 0.13) to (NPm2)-G (**Fig. 3B**). These data suggest that NP exoN activity is essential for arenavirus infectivity in IFN-competent cells and that JUNV NP exoN provided as a transgene can partially compensate for the deficiency of PICV NP exoN mutants during single-cycle replication.

**Figure 3.**
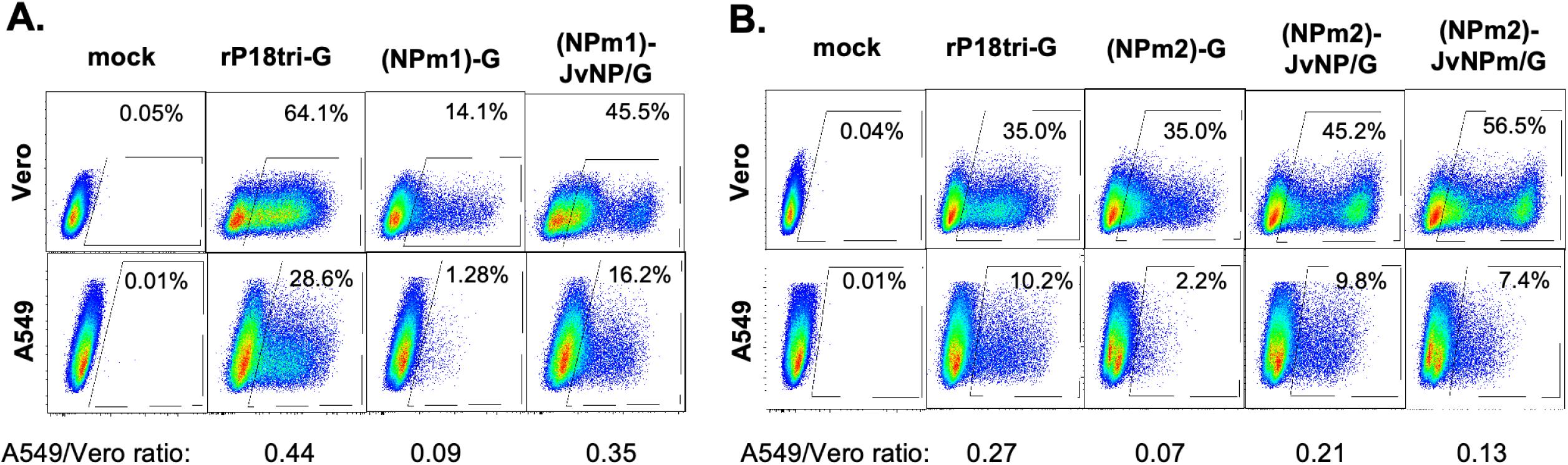
Expression of WT JUNV NP, but not exoN-deficient NP, as a transgene rescued single-cycle replication of exoN-deficient rP18tri-G mutants in A549 cells. Vero and A549 cells were infected with mock or the indicated viruses at an MOI of 0.3 for 16 h, each in triplicate. (**A**) Representative FACS plots showing GFP expression and the average percentage of GFP-positive cells for the set of (NPm1)-G viruses with or without JUNV NP transgene, along with mock-infected cells and the rP18tri-G control. (**B**) Representative FACS plots for the set of (NPm2)-G viruses expressing either WT or exoN-deficient JUNV NP, along with mock-infected cells and the rP18tri-G control. The average percentage of GFP-positive cells in A549 and Vero cells was used to calculate the A549/Vero ratio for each virus, shown at the bottom.

### JUNV NP exoN activity strongly inhibits early induction of IFN-Is

JUNV NP exoN likely contributes to arenavirus single-cycle infectivity in IFN-competent cells by blocking early induction of the IFN-I signaling. To evaluate this hypothesis, we quantified the level of selected ISGs, including PKR, RIG-I, MDA5, and ISG15, in the infected A549 cells at 16 hpi by RT-qPCR. Neither rP18tri-G nor (NPm1)-JvNP/G infection activated these genes, whereas the exoN-deficient (NPm1)-G strongly induced their expression (**Fig. 4A**). We also measured IFN-β levels released from infected A549 cells at 16 hpi and 24 hpi. As expected, rP18tri-G infection did not induce detectable IFN-β, in contrast to the highly elevated IFN-β by (NPm2)-G infection (**Fig. 4B**). (NPm2)-JvNP/G, but not (NPm2)-JvNPm/G, infection produced significantly lower IFN-β levels compared to (NPm2)-G at 16 hpi (**Fig. 4B**). Thus, functional JUNV NP exoN expressed as a transgene in a recombinant arenavirus system can suppress early induction of IFN-Is during the single-cycle infection.

**Figure 4.**
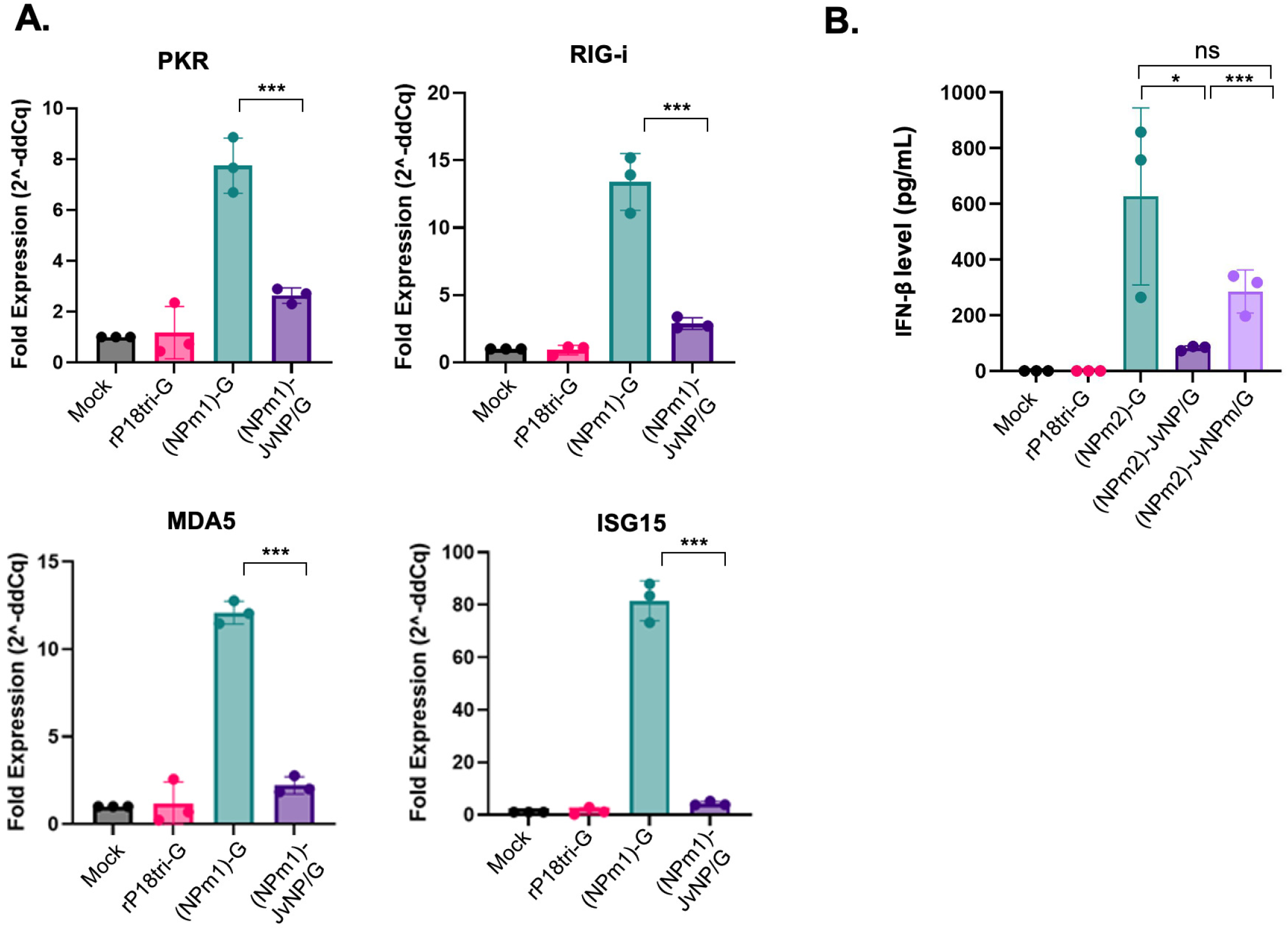
Expression of WT JUNV NP, but not exoN-deficient NP, as a transgene in the exoN-deficient rP18tri-G mutants suppressed IFN-I responses in A549 cells. A549 cells were infected with mock or the indicated viruses, each in triplicate, for 16 h. (**A**) mRNA levels of PKR, RIG-i, MDA5, and ISG15 in the infected cells were quantified by RT-qPCR and expressed as fold increase relative to mock infection for each virus. (**B**) IFN-β levels in the supernatants. ***, p < 0.001; *, p < 0.05; ns, no statistical significance.

### Quantification of infectious virus titers of recombinant reporter viruses by FACS

To accurately compare viral single-cycle replication capacity among different rP18tri mutant viruses, we need to quantify the production of infectious viruses. However, these exoN-deficient mutants produced small turbid plaques that cannot be reliably counted in the traditional plaque assay, resulting in inaccurate viral titers. As all recombinant viruses express GFP in the infected cells, we used the GFP-positive units per mL (GPU/mL) to represent the infectious virus titers and developed a FACS-based quantification assay. Viral supernatants in serial dilutions were used to infect 4x10^5^ Vero cells for 10 h, when GFP expression can be readily detected by FACS but no infectious virus has been produced (**Fig. 5A**). Within the appropriate dilution range, infectious virus titer correlates linearly with the number of GFP-positive cells. As a proof-of-concept, a viral stock of rP18tri-G, with a known PFU/mL titer of 1.24x10^6^, was quantified to be 1.44x10^7^ GPU/ml (**Fig. 5B**). The GPU titer is about 10-fold higher than the PFU titer for rP18tri, possibly because plaque formation requires multiple rounds of replication from a single infectious particle, whereas GFP expression reflects only successful viral entry and gene expression. Using this assay, the (NPm2)-JvNP stock was determined to have a titer of 5.70x10^5^ GPU/mL (**Fig. 5B**).

**Figure 5.**
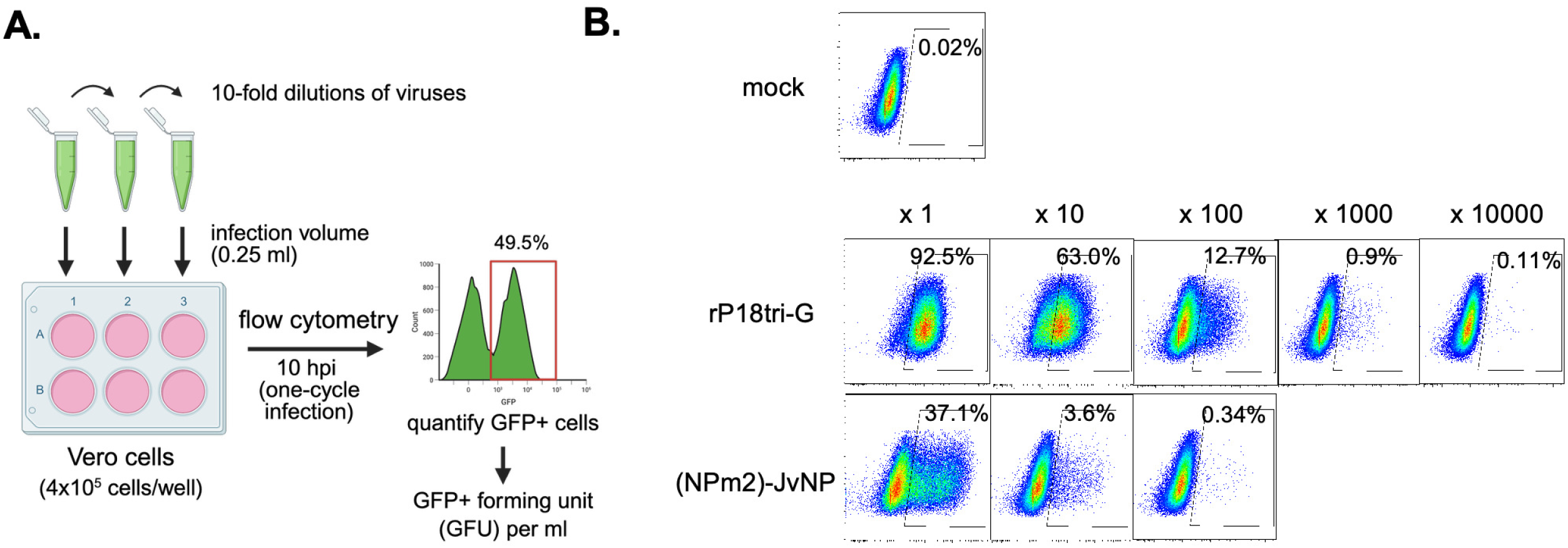
Quantification of infectious reporter virus titers by FACS based on GFP-positive units. (**A**) Schematic illustration of the procedure used to quantify GFP-positive units (GPU) per ml. Viral samples are serially diluted, 250 μL of each dilution is used to infect 4x10^6^ Vero cells. At 10 hpi, GFP-positive cells were quantified by FACS and the counts from appropriate dilutions were used to calculate GPU/ml. (**B**) Representative FACS plots showing GFP expression in infected Vero cells at 10 hpi for mock, rP18tri-G, and (NPm2)-JvNP samples at the indicated dilutions. The appropriate dilution range used to calculate GPU/ml was 100 - 10,000 for rP18tri-G and 1 - 100 for (NPm2)-JvNP.

### NP exoN activity is required for infectious virus particle production from the IFN-competent A549 cells

Using the above-described GPU assay, we measured infectious viral particle production from Vero and A549 cells at 16 hpi after single-cycle replication. Results from triplicate samples are shown as the mean ± standard deviation for the (NPm1)-G mutants (**Table 1**) and (NPm2)-G mutants (**Table 2**) in both cell lines, along with the calculated A549-to-Vero (A/V) ratio, which indicates relative viral production in IFN-competent cells. A similar trend was observed for both sets of mutants. Compared to the control rP18tri-G virus, which produced similar levels of infectious particles in Vero and A549 cells, the exoN-deficient (NPm1)-G and (NPm2)-G viruses generated significantly fewer infectious viruses in A549 cells than in Vero cells, resulting in a markedly lower A/V ratio. Expression of WT JUNV NP in the (NPm1)-G or (NPm2)-G backbone increased infectious particle production in A549 cells, as reflected by a higher A/V ratio, whereas expression of the exoN-deficient JUNV NP mutant in the (NPm2)-JvNPm/G virus did not. These data suggest that the NP exoN activity is important for infectious viral particle formation in IFN-competent cells.

**Table 1.** Infectious virus titers (GPU/ml) produced from cells after single-cycle infection (16 hpi) with WT and (NPm1)-G mutants.

| Virus | A549 | Vero | A549/Vero (A/V) ratio |
| --- | --- | --- | --- |
| **rP18tri-G** | $1.08 \times 10^5 \pm 1.55 \times 10^4$ | $1.81 \times 10^4 \pm 2.64 \times 10^3$ | 5.98 |
| **(NPm1)-G** | $2.93 \times 10^2 \pm 1.63 \times 10^2$ | $1.35 \times 10^3 \pm 3.32 \times 10^1$ | 0.22 |
| **(NPm1)-JvNP/G** | $5.50 \times 10^3 \pm 2.17 \times 10^2$ | $4.40 \times 10^3 \pm 9.89 \times 10^2$ | 1.25 |

**Table 2.** Infectious virus titers (GPU/ml) produced from cells after single-cycle infection (16 hpi) with WT and (NPm2)-G mutants.

| Virus | A549 | Vero | A549/Vero (A/V) ratio |
| --- | --- | --- | --- |
| **rP18tri-G** | $1.67 \times 10^5 \pm 9.29 \times 10^3$ | $5.33 \times 10^4 \pm 1.00 \times 10^4$ | 3.13 |
| **(NPm2)-G** | $2.03 \times 10^3 \pm 2.44 \times 10^2$ | $3.15 \times 10^4 \pm 6.36 \times 10^3$ | 0.06 |
| **(NPm2)-JvNP/G** | $3.81 \times 10^4 \pm 5.33 \times 10^3$ | $9.89 \times 10^4 \pm 6.48 \times 10^3$ | 0.38 |
| **(NPm2)-JvNPm/G** | $4.21 \times 10^3 \pm 4.61 \times 10^2$ | $7.98 \times 10^4 \pm 1.66 \times 10^4$ | 0.05 |

To assess whether the reduced production of infectious particles was linked to decreased virion RNA levels, we extracted virion RNAs from the released viral particles in the same supernatants described above and quantified PICV NP RNA by RT-qPCR. The NP RNA copy numbers per ml are shown as the mean ± standard deviation from triplicate samples for the (NPm1)-G mutants (**Table 3**) and (NPm2)-G mutants (**Table 4**). Surprisingly, all viruses, including (NPm1)-G and (NPm2)-G, produced comparable levels of virion RNAs in both A549 and Vero cells, resulting in similar A/V RNA ratios. These data suggest that the NP exoN activity is not necessarily required for virion RNA incorporation into viral particles.

**Table 3.** Virion RNA levels (NP RNA copy number per ml) in the supernatants after single-cycle infection (16 hpi) with WT and (NPm1)-G mutants.

| Virus | A549 | Vero | A549/Vero (A/V) ratio |
| --- | --- | --- | --- |
| **rP18tri-G** | $1.02 \times 10^6 \pm 8.62 \times 10^4$ | $7.02 \times 10^5 \pm 4.00 \times 10^4$ | 1.45 |
| **(NPm1)-G** | $6.25 \times 10^4 \pm 1.42 \times 10^4$ | $6.51 \times 10^4 \pm 2.38 \times 10^4$ | 0.96 |
| **(NPm1)-JvNP/G** | $1.93 \times 10^5 \pm 2.62 \times 10^5$ | $1.92 \times 10^5 \pm 5.96 \times 10^4$ | 1.00 |

**Table 4.**
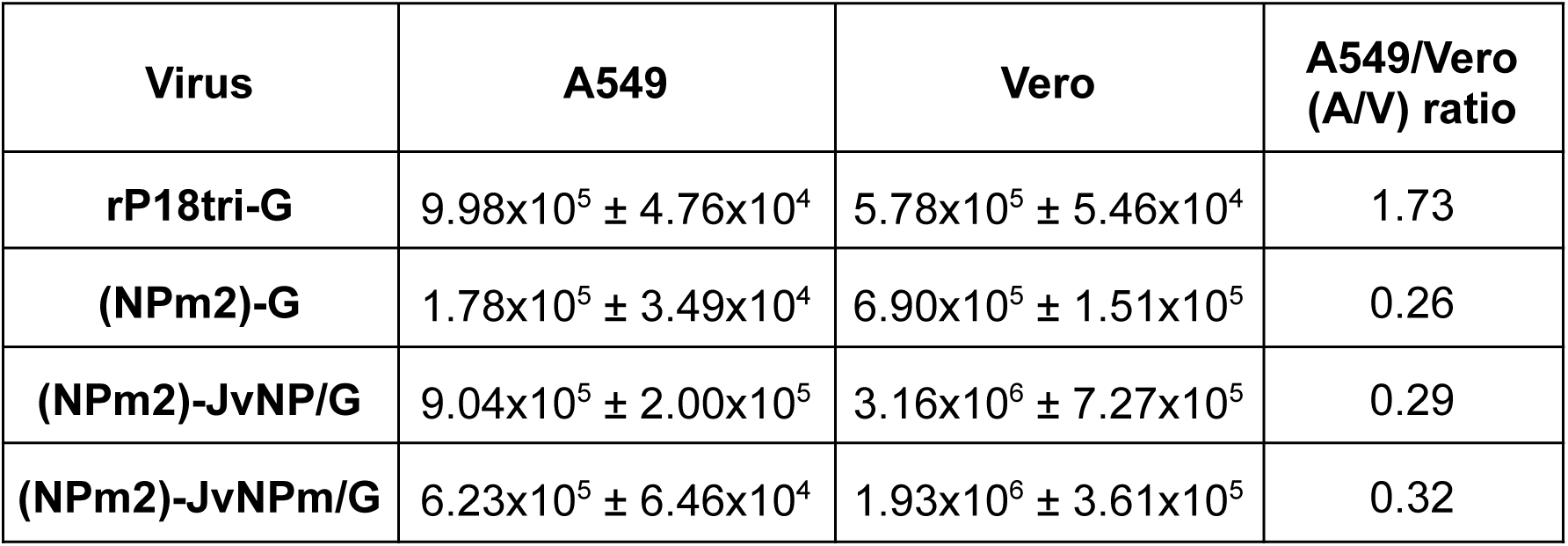
Virion RNA levels (NP RNA copy number per ml) in the supernatants after single-cycle infection (16 hpi) with WT and (NPm2)-G mutants.

We therefore calculated the viral RNA copy number per infectious particle (R/V) for each virus in Vero and A549 cells, and normalized the values to those of the control rP18tri-G virus (**Table 5**). In Vero cells, all mutant viruses showed R/V ratios similar to rP18tri-G, with normalized R/V ratios ranging from 1.0 to 3.1. In contrast, substantial differences were observed in A549 cells: the NP exoN-deficient (NPm1)-G and (NPm2)-G viruses exhibited R/V ratios 20–30-fold higher than rP18tri-G. Expression of WT JUNV NP as a transgene in the (NPm1)-JvNP/G and (NPm2)-JvNP/G viruses significantly reduced the R/V ratio to ∼4-fold above rP18tri-G levels, whereas expression of an exoN-deficient JUNV NP in the (NPm2)-JvNPm/G virus failed to lower the ratio, remaining ∼35-fold higher than rP18tri-G. These findings suggest that loss of NP exoN activity leads to the accumulation of defective virion RNAs in IFN-competent cells, supporting a critical role of NP exoN in producing infectious particles by ensuring the generation of functional virion RNAs.

**Table 5.**
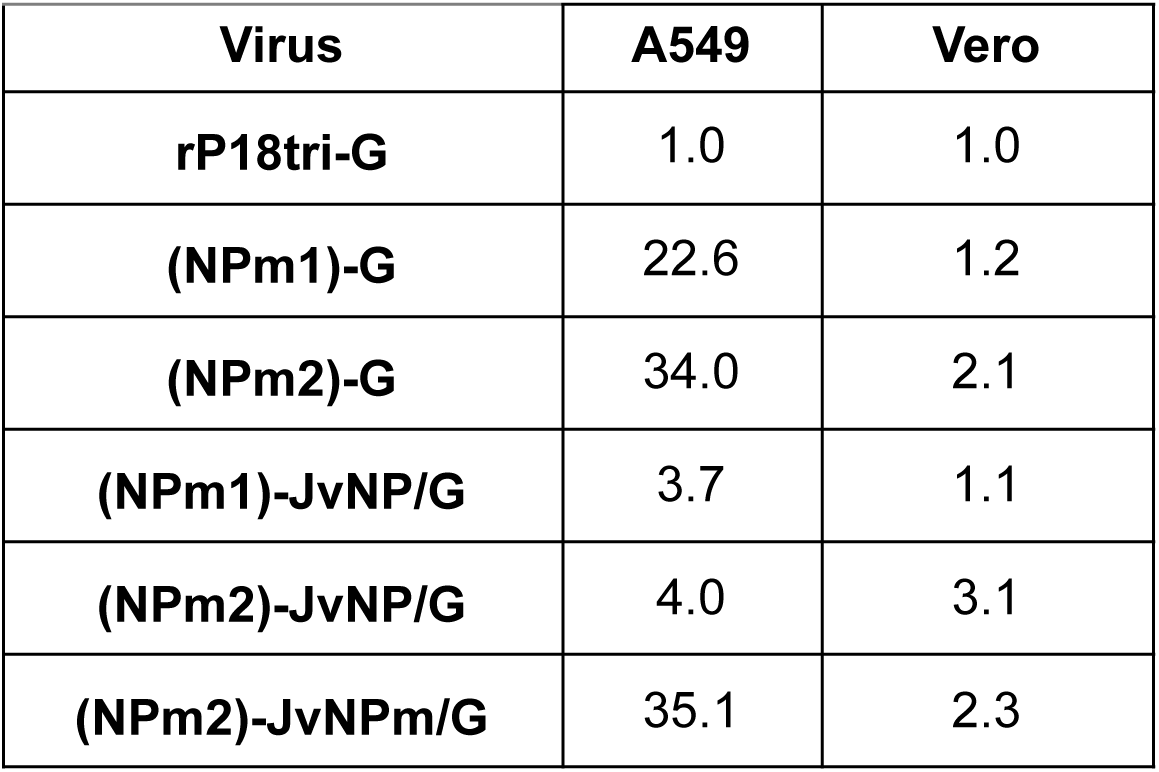
Normalized viral RNA copy number per infectious virus (R/V).

## DISCUSSION

In this study, we analyzed the functional role of arenaviral NP exoN activity during the viral infection cycle using our safe and versatile rP18tri viral vector system. We introduced mutations to abolish the PICV NP exoN activity in the rP18tri virus backbone; the resulting rP18tri-based (NPm1)-G and (NPm2)-G reporter viruses strongly activated IFN-I responses and failed to replicate efficiently in the IFN-competent A549 cells. Expression of WT JUNV NP as a transgene in the (NPm1)-G and (NPm2)-G virus backbones did not rescue multi-cycle replication but markedly increased viral infectivity with suppressed IFN-I induction during single-cycle replication, in contrast to an exoN-deficient JUNV NP mutant. Further characterization of these recombinant viruses in IFN-competent A549 cells after single-cycle replication revealed substantially increased levels of replication-defective virion particles in the absence of NP exoN activity.

Our study provides strong evidence that arenavirus NP exoN activity suppresses IFN-I induction during viral infection. Previous studies using overexpression of various arenavirus NP proteins in transfected cells (36, 37), together with structural analyses of several NP proteins (14, 15, 36), have demonstrated that NP suppresses IFN-I activation through its conserved exoN domain located in the C-terminal region. The essential role of NP exoN in IFN-I suppression during infection has also been shown using recombinant exoN-deficient PICV and LASV mutants (19–21). However, NW arenavirus pathogens such as JUNV have been reported to strongly induce IFN-I production in infected cells (38), raising the question of whether JUNV NP, despite containing a conserved exoN domain, can inhibit IFN-I induction during authentic viral infection. Using a trisegmented PICV (rP18tri) vector system, we demonstrated that NP exoN-dependent IFN-I suppression is critical for productive viral replication (**Figs. 1 and 4**). Furthermore, expression of JUNV NP, but not its exoN-deficient mutant, as a transgene in the rP18tri(NPm)-G virus backbone significantly reduced IFN-I induction during single-cycle replication (**Fig. 4**). Thus, similar to other arenavirus NP proteins, JUNV NP has a functional exoN domain that is capable of suppressing the IFN-I induction upon arenaviral infection. Nevertheless, JUNV NP could not restore multi-cycle replication of exoN-deficient rP18tri(NPm)-G viruses in A549 cells (**Fig. 2**), and several explanations are possible. Mateer et al. previously reported that JUNV NP does not degrade dsRNA to the same level as LASV NP in respective virus-infected cells (20). Thus, JUNV NP may be less effective in IFN-I suppression compared to other arenavirus NP proteins and cannot fully compensate for PICV NP exoN activity. This hypothesis could be tested by comparing JUNV NP, LASV NP, and PICV NP in the same rP18tri(NPm)-G virus backbone in future studies. Alternatively, a more likely explanation is that a significant proportion of tri-segmented rP18tri(NPm)-JvNP/G viral particles fail to package a complete genome during multi-cycle replication, and that the absence of JUNV-NP-encoding S1 segment leads to insufficient IFN-I suppression. Further studies are needed to assess the efficiency of virion RNA packaging for bi- and tri-segmented rPICVs and to determine whether loss of an RNA segment encoding a functional NP exoN activity correlates with heightened IFN-I responses. Finally, it should be noted that the trisegmented rP18tri virus differs from authentic JUNV in infectivity, and the precise biological role of JUNV NP in modulating IFN-I responses will require studies using a recombinant JUNV system.

This study reveals a novel biological role of arenaviral NP exoN in promoting the production of infectious viral particles in IFN-competent cells. We found that recombinant viruses lacking exoN activity produced substantially higher levels of defective virion particles in IFN-competent A549 cells compared with IFN-deficient Vero cells. This defect can be rescued by expression of WT JUNV NP, but not by an exoN-deficient JUNV NP mutant, provided as a transgene. While NP exoN has been known to be essential for arenavirus replication in IFN-competent cells, this is, to our best knowledge, the first demonstration of its key role in generating infectious viral particles from these cells. Similar to other segmented RNA viruses, only a fraction of arenavirus particles are deemed to be infectious. Arenaviral quasispecies with genetic variants at essential sites, defective viral genomes (DVGs) with large deletions, and failure to package one or more RNA segments can all lead to defective particles (39, 40). NP exoN may promote the formation of infectious particles through direct and/or indirect mechanisms. IFN-I activation may increase the generation of defective virion particles by elevating error rates during viral RNA replication and/or disrupting viral RNA packaging. Thus, NP exoN may indirectly enhance infectious particle production by suppressing IFN-I responses. NP exoN may also directly improve the fidelity of viral RNA polymerase during genomic RNA synthesis. Notably, coronavirus-encoded exoN, nsp14, has proofreading activity during viral RNA synthesis (41–43). A recent study using a recombinant exoN-deficient LASV mutant suggested that LASV NP exoN may also play a role in proofreading (44). However, most of these studies demonstrating the proof-reading activity of viral exoN were conducted in IFN-deficient cells, whereas our findings of arenaviral NP exoN-dependent production of infectious viral genomes were observed in IFN-competent cells. Further studies are needed to determine the exact mechanism(s) by which NP exoN contributes to infectious particle formation, particularly in IFN-competent cells.

In summary, our study identifies a novel role for arenavirus NP exoN in the production of infectious particles in IFN-competent cells, confirms its established function in suppressing IFN-I responses, and demonstrates the utility of the rP18tri vector system for evaluating the biological functions of a transgene, such as NP in this study, during authentic arenavirus infection. This research further emphasizes the importance of the arenaviral NP exoN as a promising target for antiviral development.

## ACKNOWLEDGMENTS

We thank K.K. Conzelmann for providing BSRT7-5 cells. This work was supported in part by NIH award R01AI131586 to H.L. and Y.L.

